# Host protein CD63 enhances viral RNA replication by interacting with human astrovirus nonstructural protein nsP1a/4

**DOI:** 10.1101/331835

**Authors:** wei zhao, Nian liu, Xiao Li Tao, Chun hong Zheng, Xiang yu Li, Man Yu, Yong gang Li

**Affiliations:** College of Basic Medical Sciences, Jinzhou Medical University, Jinzhou 121200, Liaoning, China; Biological Anthropology institute, Jinzhou Medical University, Jinzhou 121200, Liaoning, China

**Author notes:** Corresponding author: College of Basic Medical Sciences, Jinzhou Medical University, No. 40, the Third Section of SongPo Rd., Jinzhou City 121200, Liaoning Province, China.

**Keywords:** human astrovirus, HAstV, protein-protein interaction, nsP1a, nsP1a/4, CD63, virus replication

## Abstract

Human astrovirus nonstructural protein nsP1a/4, located at the C-terminal end of nsP1a, is thought to be involved in the regulation of RNA replication and capsid maturation;however, its rolesviral growth and virulence are not well understood. We investigated the intracellular host proteins that interact with nsP1a and explored the potential roles of the interaction in the pathogenesis of human astrovirus infection. We screened 14 independent proteins with a cDNA library derived from Caco-2 cells using a yeast two-hybrid technique. Deletion analysis revealed that interaction between the nsP1a/4 domain and the large extracellular loop (LEL) domain of the human protein CD63 is necessary for astrovirus replication. The interaction was confirmed by glutathione-S-transferase (GST) pull-down assays and co-immunoprecipitation assays. Confocal microscopy showed that nsP1a/4 and CD63 co-localized in the cytoplasm of infected cells. Over expression of CD63 promoted viral RNA synthesis, whereas knockdown of CD63 markedly decreased viral RNA levels. Those results suggest that CD63 plays a critical role in human astrovirus RNA replication. The interaction between CD63 and nsP1a/4 provides a channel to further understand the roles of interactions between host and virus proteins in astrovirus infection and release.

**IMPORTANCE:** Human astroviruses cause gastroenteritis in young children and immunocompromised patients. In this study, we provide evidence that nsP1a/4, a nonstructural protein located at the C-terminal end of the human astrovirus nsP1a polyprotein, interacts with the host protein CD63. Over expression of CD63 promoted viral RNA replication, whereas knockdown of CD63 decreased virus RNA replication, indicating that CD63 plays a critical role in the human astrovirus life cycle.

---

Viruses are obligate intracellular pathogens thatutilize host-cell machinery to infect cells, replicate themselves, and then exit the cells for another round of infection. Human astroviruses (HAstVs) are a group of non-enveloped, single-stranded, and positive-sense RNA viruses in the family *Astroviridae*. HAstVs are causative agents of viral diarrhea in young children, immunocompromised patients, and elderly individuals (1-5). The astrovirus genome has three open reading frames (ORFs); ORF1a and ORF1b encode two nonstructural polyproteins (nsPs), and ORF2 encodes the capsid protein (6). The nsP1a polyprotein, transcribed from ORF1a, is involved in viral transcription and replication. When HAstV infects a host cell, nsP1a is cleaved into at least four products, named nsP1a/1, nsP1a/2, nsP1a/3 (protease), and nsP1a/4. The nsP1a/4 protein is located at the C-terminal end of nsP1a.Several domains have been identified in nsP1a/4, including two coiled-coil regions, a death domain (DD), a nuclear localization signal (NLS), a putative viral genome-linked protein(VPg), and a hypervariable region (HVR)(7-12). It was confirmed that nsP1a/4 plays important roles in the activation of capsid maturation and the regulation of RNA replication (13, 14). To better understand the roles of nsP1a and nsP1a/4 in HAstV infection and replication, we characterized host proteins that interact with nsP1a or nsP1a/4 by utilizing a yeast two-hybrid approach. We identified at least 14 host proteins that could bind to nsP1a, one of which was CD63.

CD63, a member of the tetraspan transmembrane protein family, is a four-span membrane protein that is widely distributed in multicellular organisms and known to be associated with virus functions, such as adhesion, fusion, and trafficking (15, 16). Tetraspanins, a large superfamily of cell-surface membrane proteins characterized by four transmembrane domains, have the unique property to form a network of protein-protein interactions through associations with multiple membrane proteins that are involved in several infectious diseases (17, 18). There is increasing evidence suggesting that intracellular pathogens, especially viruses, “hijack” tetraspanins when entering, traversing and exiting cells during the course of infection. Several studies have reported that tetraspanins, such as CD151 (19), CD81 (20, 21), CD82 (22), CD9 (23), and CD63 (24), are key players in the lifecycles of many viruses.

In this study, we used co-immunoprecipitation and glutathione-S-transferase(GST) pull-down assaysto confirm that CD63 can interact with nsP1a/4.Confocal microscopy revealed that the CD63 large extracellular loop (LEL) domain co-localizes with nsP1a/4 in the cytoplasm. CD63 knockdown or overexpression strongly affected the replication of HAstV in Caco-2 cells. Those findings indicate that CD63 plays an important role in the HAstVlife cycle.

## RESULTS

### A yeast two-hybrid screen withnsP1aand a Caco-2 cDNA library identified CD63 as a host protein that interacts with nsP1a

Weinvestigated the interactionsbetween HAstV nsP1a and host proteins using a protein-protein overlay assay. We amplified thensP1a gene by PCR and inserted the resulting PCR product (2781bp) by recombination cloning into the yeast two-hybrid (Y2H) bait vector pGBKT7 as a C-terminal fusion with a Gal4 DNA-binding domain (BD). We confirmed the correct constructs by enzyme digestion and DNA sequence analysis (Fig 1A). When introduced alone into yeast Y2HGold cells, the Gal4 (BD)-nsP1a bait construct wasnot toxic to the yeast and did not auto-activate the Y2H reporter gene, indicating that the construct was suitable for use in a Y2H screen (Fig 1B).

**Fig 1.**
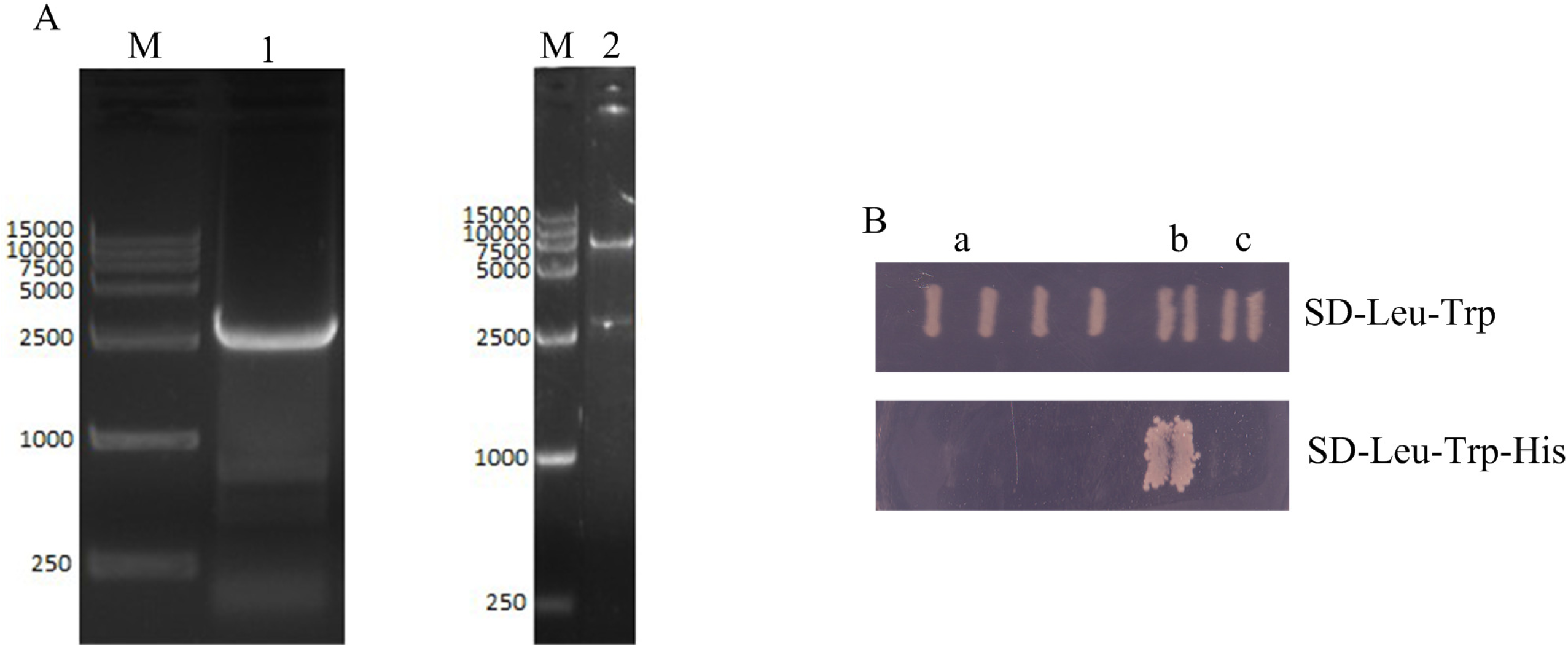
Construction of nsP1a bait plasmid and detection of auto-activation. (A) Construction of pGBKT7-nsP1a bait plasmid: Lane M: DL15000 DNA marker; Lane 1: amplified fragment of nsP1a; Lane 2: confirmation of pGBKT7-nsP1a by digestion with BamH I and Pst I. (B) Determination of the auto-activation of the pGBKT7-nsP1a bait plasmid in yeast cells. The pGBKT7-nsP1a bait and pGBKT7 plasmids were used to transform Y2HGold cells and then grown on different plates. Co-transformants containing pGADT7-T and pGBKT7-53 were grown on SD-Leu-Trp-His plates as a positive control. Co-transformants containing pGADT7-Lam and pGBKT7-T were grown on SD-Leu-Trp-His plates as negative control. a: pGBKT7-ORF1a +pGADT7; b: pGBKT7-53 + pGADT7-T;c: pGBKT7-Lam + pGADT7-T.

Using a Y2H assay with Gal4 (BD)-nsP1a as bait and a human Caco-2 cDNA prey library, we screened approximately 2.5×10^5^clones and identified 14 positive bait-prey interactions. Sequence and bioinformatics analysis of the 14 positive prey plasmids,listed by their initial positive-interaction identification number,indicated that they represented 14 different human cDNAs(data not shown). We next measured the strength of interaction between thensP1a bait and each of the 14 human prey proteins in yeast using a qualitative growth assay and quantitative β-galactosidase activity assays. We performed bioinformatic analysis of the human proteins confirmed by the qualitative growth assay to gain functional insights into the potential interactions between those proteins and nsP1a (Fig 2). We selected one host protein, identified as tetraspan transmembrane protein CD63 (GenBank accession number: NM_001040034.1), for further study.

**Fig 2.**
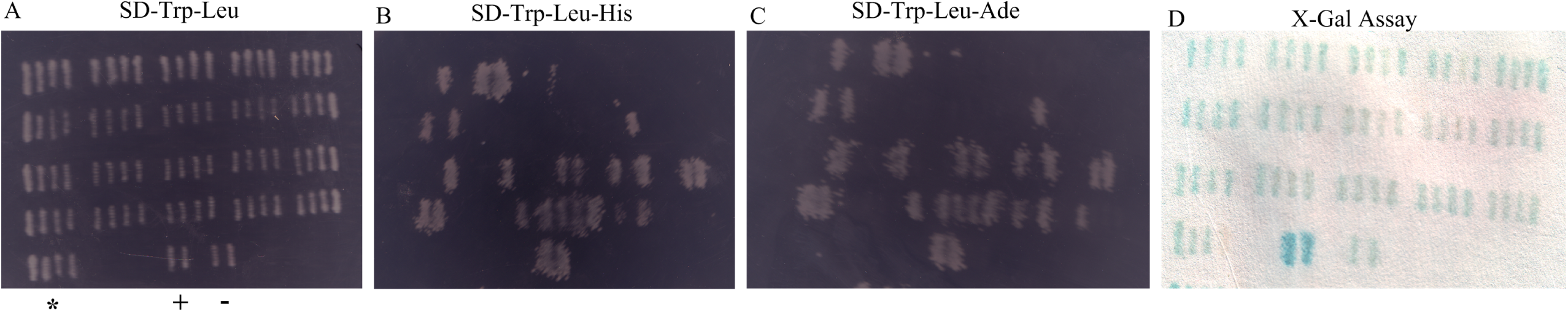
Analysis of putatively positive colonies of Y2HGold cells. Y2HGold clones containingthe cDNA library were grown on dropout medium lacking tryptophan and leucine (SD-Trp-Leu). There were 25 clones, and the total number cloned was 2.5×10^5^.Yeast cells containing the bait (HAstV nsP1a) and prey plasmids (Caco-2 cell cDNA library) werescreened for interaction on synthetic dropout media. (A) Screen for interaction on SD-Trp-Leu medium. (B) Screen for interaction on synthetic dropout medium lacking tryptophan, leucine,and histidine(SD-Trp-Leu-His). (C) Screen for interaction on synthetic dropout medium lacking tryptophan, leucine, histidine, and adenine (SD-Trp-Leu-His-Ade). (D)Confirmation of positive interactions on SD-Trp-Leu-X-Gal medium; 15 clones demonstrating protein interactions between HAstV and nsP1a were obtained. Co-transformation with pGADT7-T and pGBKT7-Lam was used as a negative control (-).Co-transformation with pGADT7-T and pGBKT7-53 was used as a positive control (+). Co-transformation with pGBKT7-nsP1a and pGADT7 was used as an auto-activation control (*).

### nsP1a interacts with CD63 *in vivo*

We transfected HEK293 T cells with PEF-HA-nsP1a and pcDNA3.1-3flag-CD63.After 48 h, we performed immunoprecipitation with the cell lysates from those cells using an anti-Flag M2 Affinity Gel (Sigma, St. Louis, MO, USA). We analyzed the resulting protein complexes by western blot with anti-Flag or anti-HA antibody (Fig. 3A). We detected no proteins in control lysates from cells transfected with empty vector (Fig. 3A, lane 1).We detected CD63 protein only in the lysates from cells that wre transfected with pcDNA3.1-3flag-CD63 (Fig. 3A, lane 2).Likewise, we detectednsP1a protein only in the lysates from cells that were transfected with PEF-HA-nsP1a (Fig. 3A, lane 3). Both nsP1a and CD63 were readily detected in the lysates from cellsthat were co-transfected with pcDNA3.1-3flag-CD63 and PEF-HA-nsP1a. The HA antibody was able to pull down CD63-Flag together with nsP1a-HA (Fig. 3A, lane4). Likewise, the Flag antibody was able to pull down nsP1a-HA together with CD63-Flag (Fig. 3A, lane4). Taken together, the results suggested that nsP1a was able to interact with the CD63 *in vivo*.

### nsP1a interacts with CD63 *in vitro*

To further investigate the interaction between nsP1a and CD63, we determined whether the interaction occurs *invitro.* We expressed recombinant full-length nsP1a-GST fusion protein (≈118kDa) induced by isopropyl β-D-thiogalactopyranoside(IPTG)in *Escherichia coli* (Fig. 3B lanes 1–6). We also detected purified nsP1afrom the bacteria (Fig. 3B lane7) and human CD63 His Tag commercialized recombinant protein (≈26kDa) by western blot (Fig 3B lane8). We tested the interaction between CD63 and nsP1a by GSTpull-down analysis. We immobilized nsP1a-GST on glutathione agarose and added CD63-His to assess protein-protein binding. We used an untreated gel as a negative control. We analyzed the protein in the complexes by western blot with anti-GST or anti-His antibody. Anti-GST MAb pulled down nsP1a together with CD63 (Fig. 3C). Consistent with those results, anti-His MAb could pull down CD63 protein together with nsP1a protein. In contrast, the negative control could not pull down any viral proteins. The results indicated that nsP1a could interact with CD63 *in vitro*.

**Fig 3.**
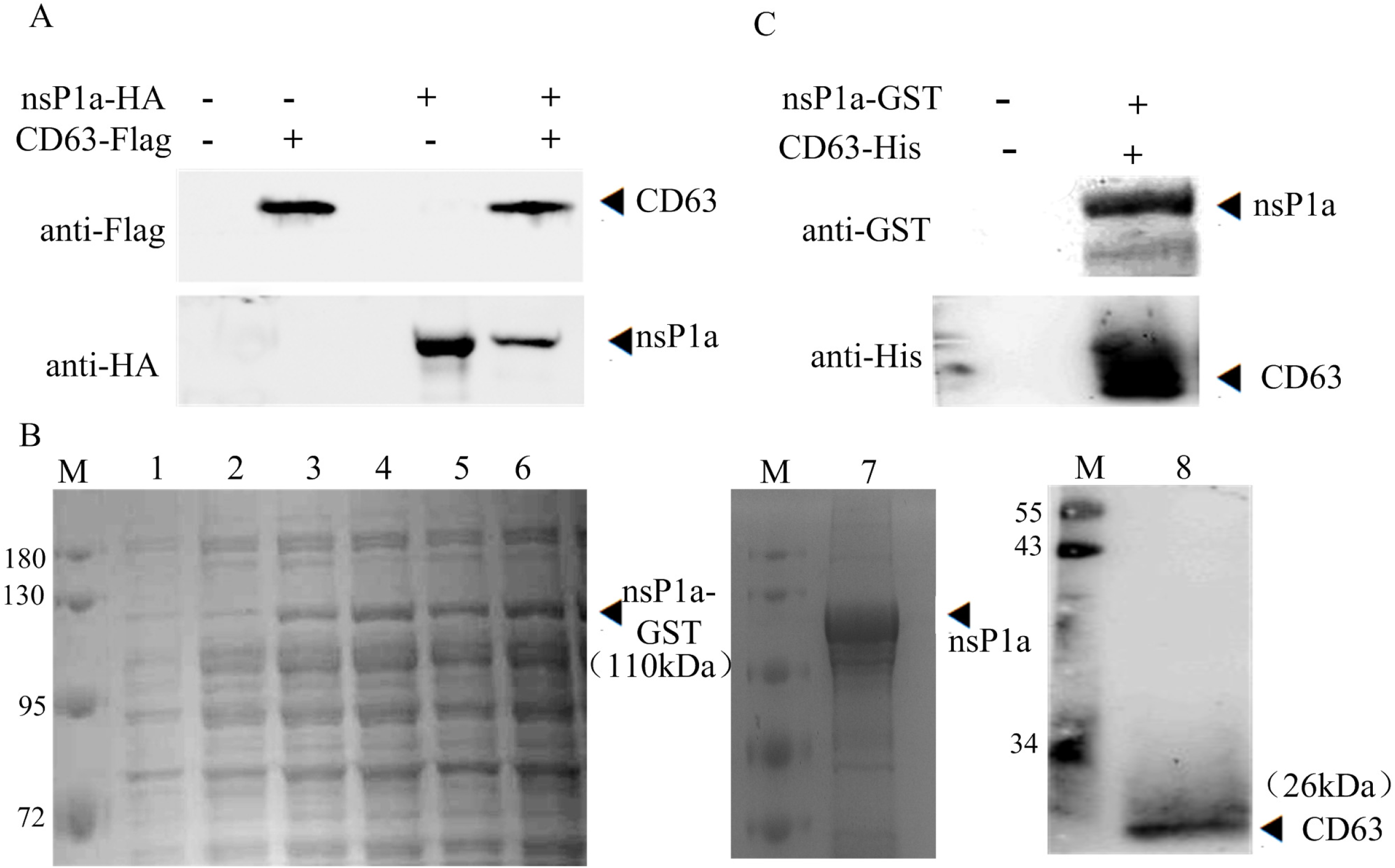
nsP1a interacts with CD63 *in vivo* and *in vitro*. (A) HEK293T cells were co-transfected with nsP1a-HA plasmid and CD63-Flag plasmid for 48 h and harvested. Cell lysates from co-transfected and untransfected control cells were immunoprecipitated with antibody against Flag or HA and then analyzedby western blot. (B) The recombinant plasmids nsP1a-GST were transfected into *E. coli* BL21 (DE3) cells. The cells were then mixed with IPTG at concentrations of 0, 0.1, 0.2, 0.4, 0.8, and1mM (lanes 1–6, respectively) to induce the expression of nsP1a.The cells were then incubated for an additional 5 h at 25°C. The purified nsP1a-GST protein (110kDa) was analyzed by SDS-PAGE and stained with Coomassie blue R-250 (lane 7). The recombinant protein CD63-His was detected by western blot using CD63 antibody. Lane M: protein molecular weight marker. (C) nsP1a-GST was immobilized on glutathione agarose. CD63-His was then added to assay protein binding. The proteins were washed, eluted from the agarose beads, and confirmed by immunoblotting. The expression of nsP1a was detected by anti-GST mAb. CD63 was detected by anti-His.

### The nsP1a/4 protein interacts with the CD63 large-loop domain

We performed binding experiments to identify the region of the nsP1a protein that interacts with CD63 *in vivo* and *in vitro.* CD63 contains 204–355 amino acids (20–30 kDa), comprising four transmembrane domains, a short extracellular loop (EC1), a long extracellular loop (EC2), a very short intracellular loop, and cytoplasmic N-terminal and C-terminal tails (Figure 4A). EC2, also called the large loop (LEL, 99aa), contains a variable region for specific protein-protein interactions. We constructed and confirmed the eukaryotic expression vector pcDNA3.1-3flag-CD63-LEL and the prokaryotic expression vector pET-29a-LEL. We expressed IPTG-induced recombinant CD63-LEL-His fusion protein in *E. coli* and recovered and purified the recombinant protein from the bacteria (Fig 4C).

**Fig 4.**
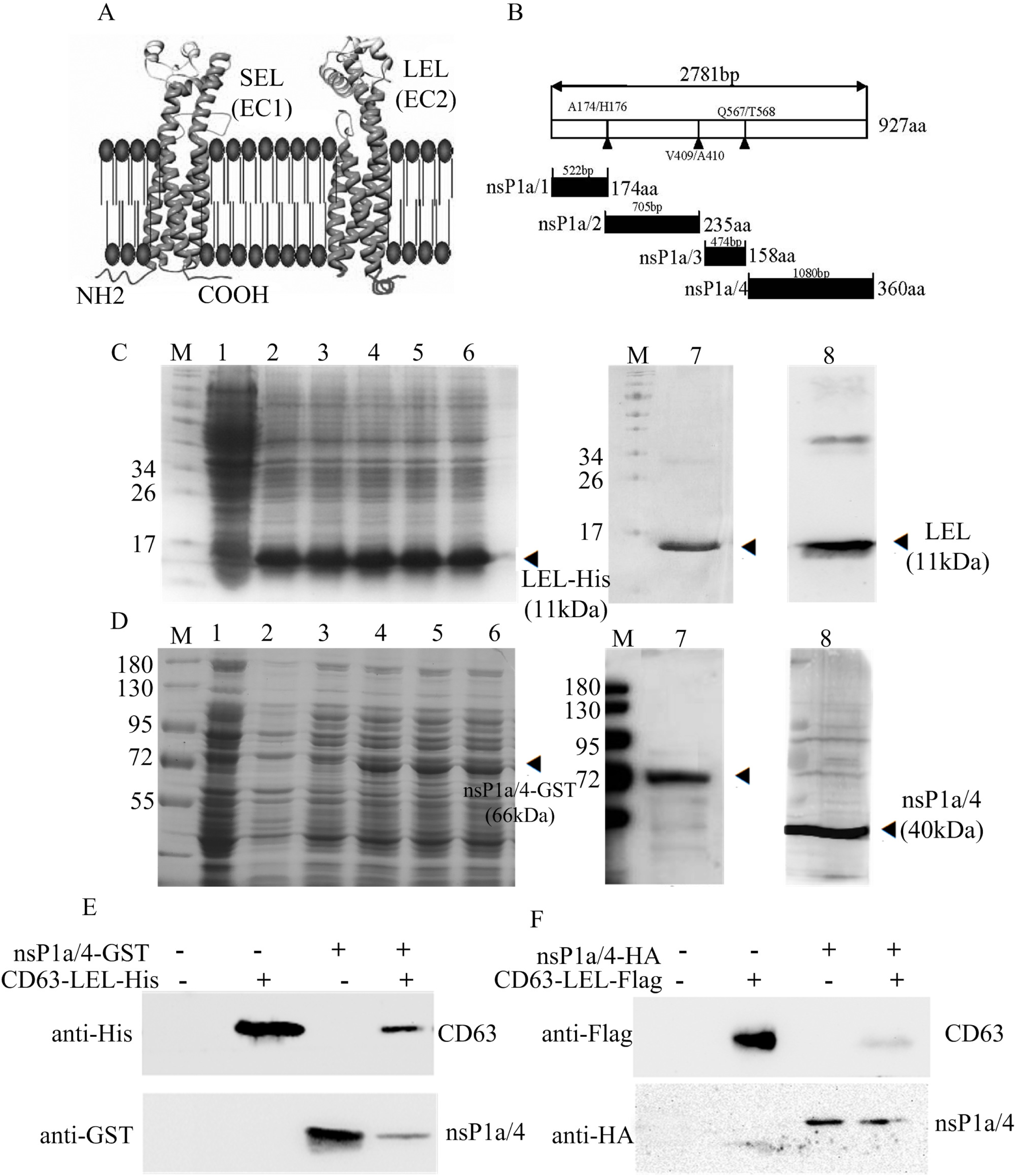
The C-terminal nonstructural protein nsP1a/4 interacts with the CD63 large loop LEL domain. Structure of the protein domains of CD63 and the LELdomain. (B)Schematic representation of the protein domains of the HAstV nsP1a protein and the four domains protein tested in this study. (C) The recombinant plasmids CD63-LEL were transfected into *E. coli* BL21 (DE3). The cells were mixed with0,0.1,0.2,0.4, 0.8, or 1mM IPTG (lanes 1–6, respectively) to induce CD63expression, and the culture was incubated for an additional 6 h at 30°C. The purified CD63-LEL-His protein (11kDa) was analyzed by SDS-PAGE and stained with Coomassie blue R-250(lane 7). The recombinant protein CD63-LEL was detected by western blot using CD63 antibody (lane 8). (D)The recombinant plasmid nsP1a/4-GST wastransfected into *E. coli* BL21 (DE3). The cells were mixed with0,0.1,0.2,0.4,0.8, or 1mM IPTG (lanes 1–6, respectively) to induce nsP1a/4expression, and the culture was incubated for an additional 6 h at 37°C. The purified nsP1a/4-GST protein (66kDa, GST26kDa) was analyzed by SDS-PAGE and stained with Coomassie blue R-250(lane 7). The recombinant protein nsP1a/4 was detected by western blot using nsP1a/4 antibody (lane 8). Lane M: protein molecular weight marker. (E) nsP1a/4-GST was immobilized on glutathione agarose. CD63-LEL-His was added to assay protein-protein binding. After washing, proteins were eluted from the agarose beads and confirmed by immunoblotting. The expression of nsP1a/4 was detected by anti-GST mAb, and CD63-LEL was detected by anti-His. (F) HEK293T cells were co-transfected nsP1a/4-HA plasmid and CD63-LEL-Flag plasmid for 48 h and harvested. Cell lysates from co-transfected and untransfected control cells were immunoprecipitated with antibody against Flag or HA followed by western blot analysis.

We amplified the four products of the nsP1a polyprotein (nsP1a/1, nsP1a/2, nsP1a/3 (protease), and nsP1a/4; Fig 4B) by RT-PCR. We used the PCR products to construct PEF-HA-1a/1, PEF-HA-1a/2, PEF-HA-1a/3, and HA-1a/4 plasmids, which we confirmed by sequencing. We expressed recombinant nsP1a/4-GST fusion protein in bacteria and isolated and purified the fusion protein from the bacteria (Fig4D). We co-transfected HEK293T cells with pcDNA3.1-3flag-CD63-LEL and either PEF-HA-1a/1, PEF-HA-1a/2, PEF-HA-1a/3, or PEF-HA-1a/4. We then performed immunoprecipitation with lysates from the transfected cells and anti-HA or anti-Flag antibodies. We identified protein-protein interactions by GST pull-down (Fig 4E) or co-immunoprecipitation (Fig 4F) assays as described above. The results showed that the CD63 LEL domain interacted with nsP1a/4 *in vivo* and *in vitro*. In contrast, no interaction between CD63-LEL and nsP1a/1, nsP1a/2, or nsP1a/3 were observed (data not shown).

### CD63-LEL co-localizes with nsP1a/4 in host cells

We used immunofluorescence and confocal microscopy to further investigate the interaction betweennsP1a/4 and CD63-LEL. We co-transfected HEK293T cells with the plasmids HA-nsP1a/4 and Flag-CD63-LEL. We then examined the subcellular localization of nsP1a/4 and CD63-LEL by confocal microscopy. The Flag-CD63-LEL (Fig. 5A) and HA-nsP1a/4 (Fig. 5B) proteins were distributed throughout the cytoplasm. We determined the position of the nucleus by DAPI staining (Fig. 5C). Statistical analysis of the images showed that the (immunofluorescence assay,IFA)signals of nsP1a/4 and CD63-LEL significantly overlapped each other (Fig 5D). That finding confirmed that nsP1a/4 protein interacts with CD63-LEL in HEK293T cells. That result along with those of a previous Y2H screen and GSTpull-down and co-immunoprecipitation assays suggested that nsP1a/4 interacts with the CD63LEL domain.

**Fig 5.**
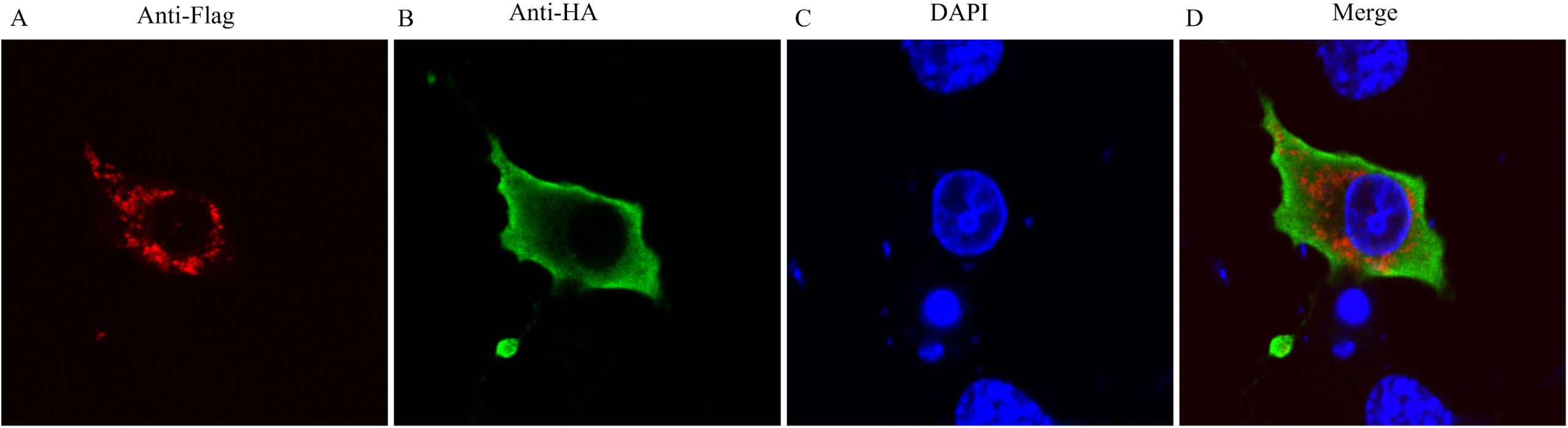
Co-localization of nsP1a/4 protein with CD63-LEL protein. 293 T cells were co-transfected with PEF-nsP1a/4-HA and pcDNA3.1(+)-CD63-LEL Flag plasmids. Cells were fixed after 48 h and subjected to indirect immunofluorescence to detect CD63-LEL-Flag (red, A) and nsP1a/4-HA (green, B) with rabbit anti-Flag and mouse anti-HA antibodies. The position of the nucleus is indicated by DAPI (blue, C) staining in the merged image. Co-expression of CD63-LEL and nsP1a/4 showing cytoplasm localization in merged images D.

### HAstV infection increases CD63 expression

HAstV-1 nsP1a has been reported to regulate the viral RNA-replication process. To determine whether HAstV-1 infection affects CD63 expression, we infected Caco-2 cells with HAstV-1 (MOI 10). We collected lysates from mock-infected (control) and HAstV-infected cells 24h and 48h after infection. We isolated total cellar RNA and quantified the expression of CD63 mRNA using real-time RT-PCR(Fig. 6A). We also performed western blot to detect CD63 protein expression in the cellular lysates (Fig 6B). The results showed that the levels of CD63 mRNA and protein 48h after infection were higher in HAstV-infected cells than inmock-infected cells, suggesting that CD63 plays an important role in HAstV-1 genome replication. Furthermore, we infected Caco-2 cells with HAstV-1(MOI 1) and measured the amount of viral RNA present in the cell culture after 24h and 48h. The results showed that the amount of viral RNA was significantly increased at 24h and 48h in the HAstV-1-infected cells compared with those in the mock-infected cells (P<0.01; Fig 6C).

**Fig 6.**
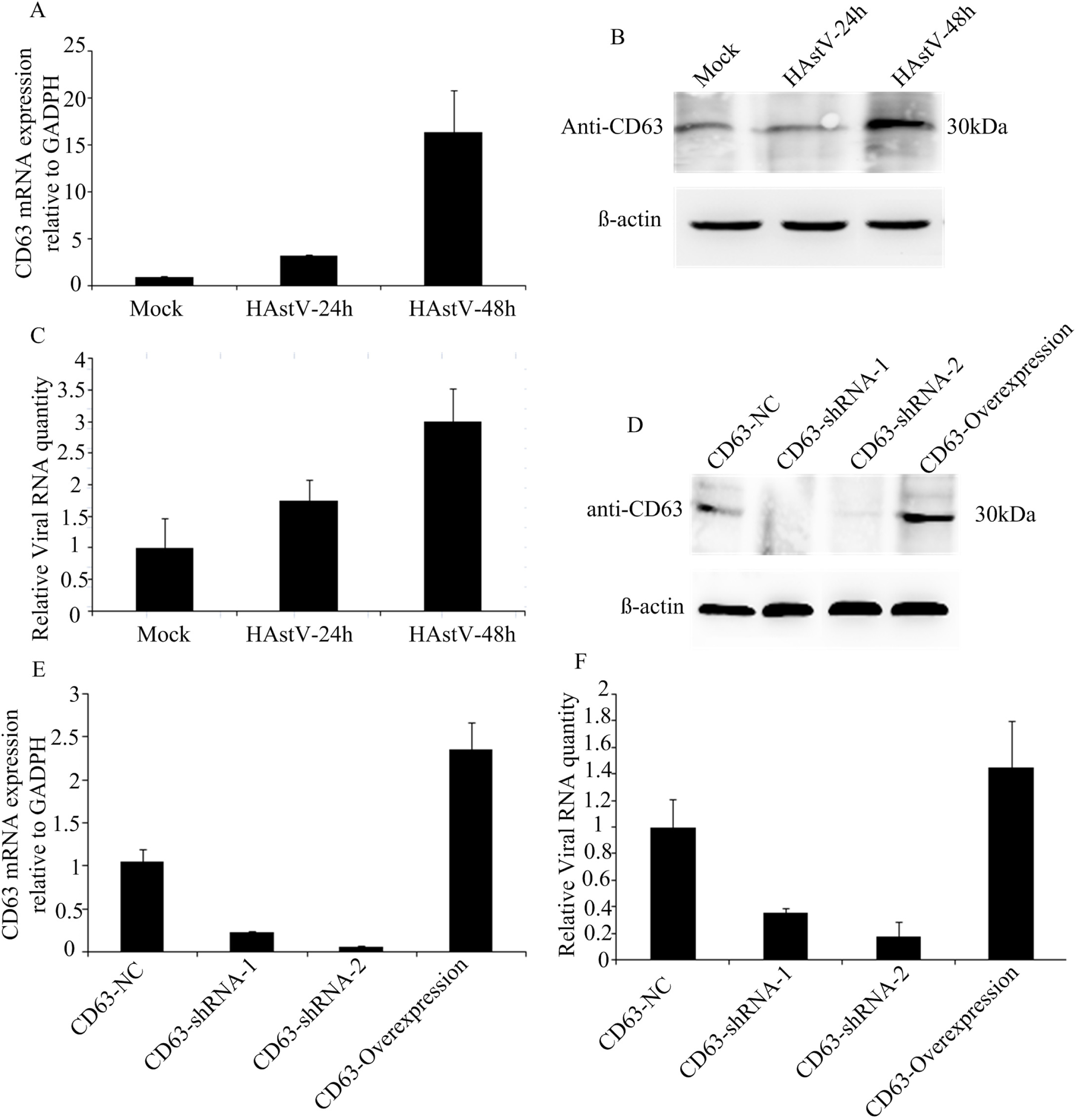
CD63 effect on HAstV-1 replication. The amount of CD63 mRNA expression in the Caco-2 cells infected with HAstV-1 virus by real-time PCR(P<0.01); (A) Real time PCR analysis of CD63 mRNA expression in Caco-2 cells infected with HAstV-1. (B) Western blot analysis of CD63 protein expression in Caco-2 cells infected with HAstV-1. (C) Intracellular viral RNA levels assayed by real-time PCR (P<0.01). (D) Overexpression of CD63 mediated by eukaryotic expression vector pcDNA3.1-3flag and knockdown of CD63 expression mediated by shRNA. (E) Intracellular CD63 mRNA expression levels assayed by real-time PCR (P<0.01). (F) Intracellular viral RNA levels assayed by real-time PCR (P<0.01).

### CD63 expression affects HAstV replication

To assess the role of CD63 in HAstV replication, we infected CD63-overexpressing and CD63-knockdown Caco-2cells with HAstV-1 and measured the level of viral RNA 48 h after infection. We assessed the overexpression and knockdown of CD63 by western blot and real-time RT-PCR, respectively (Fig. 6D and E). CD63 expression relative to the wild-type level was increased 5-fold in the cells transfected with the overexpression plasmid and decreased 10-fold in the cells transfected with the anti-CD63 shRNA.We then infected the cells with HAstV-1(MOI 1) and measured the amount of viral RNA present in the cell cultures 48h after infection. The results showed a 1.45-fold increase in the level of HAstV-1 RNA in the cultures of CD63-overexpressing cells compared with that in cultures of control cells(P<0.01). The CD63 knockdown resulted in a 6-folddecrease inintracellular viral RNA levels(P<0.01; Fig 6F). Those results indicated that CD63 plays animportant rolein the HAstV lifecycle.

## DISCUSSION

Positive-sense RNA viruses have frequently been found to replicate in association with a large number of cellular proteins (25). The C-terminal nsP1a protein of HAstV Yuc8 was shown to interact with the viral polymerase (14). Astrovirus replication and assembly have also been linked to fatty-acid synthesis, ATP biosynthesis, and cellular lipid metabolism (26), but it is unknown what host proteins contribute to a successful virus infection. We found that HAstV nsP1a or nsP1a/4 could interact with the host protein CD63 based on a Y2H screen and observations of direct interaction between nsP1a/4 and CD63 both *invivo* and *in vitro.*

Although the function of HAstVnsP1a remains unclear, nsP1a and nsP1a/4 have been suggested to be involved in many processes, including genome replication, apoptosis induction, and capsid maturation (8, 11). To better understand the role of nsP1a in viral replication, we analyzed the interaction of nsP1a with host proteins. A Y2H screen of aCaco-2 cDNA library showed that 14 independent proteins, including CD63, interacted with nsP1a.

CD63 was the first characterized tetraspanin(27). It has been shown that CD63 reacts with many different proteins either directly or indirectly, including integrins(28, 29), other tetraspanins(30,31), cell-surface receptors(32), and kinases(33). Viruses are obligate intracellular pathogens andmust utilize host-cell machinery in order to complete their lifecycle. Viruses have been shown to incorporate many host components, including tetraspan transmembrane proteins (34). It is therefore highly likely that tetraspanins play an important role in the lifecycles of viruses. We confirmed that CD63 could interact with HAstV nsP1a both *in vivo* and *in vitro*. We transfected HEK293T cells with nsP1a-HA alone or togetherwith CD63-Flag. We successfully recovered nsP1a-HA/CD63-Flag complex by co-immunoprecipitation assays of co-transfected cells. Weshowed that CD63 bound directly to nsP1a *in vitro* usingGSTpull-down analyses ofpurified recombinant GST-nsP1a protein and recombinant CD63-His protein. Those results showed that the HAstV nsP1a could interact with CD63.

In light of those results, we sought to explore the nsP1a/CD63 interaction in more detail. GSTpull-down and co-immunoprecipitation analyses showed that the nsP1a C-terminal nonstructural protein nsP1a/4 (567–926aa, 40kDa) binds to the CD63 LEL domain. The LEL of CD63 displays some conserved residues. Protein-protein interaction sites have been found in the LEL domains of other proteins (35, 36). We verified that the CD63-LEL domain interacts with nsP1a/4 *in vivo* and *in vitro*. Confocal microscopy also showed that nsP1a/4 and CD63 co-localized in the cytoplasm. Previous data indicatedthat the HAstV nsP1a polyprotein would generate at least four products. The nonstructural C-terminal protein of nsP1a, which contains ahypervariable region, has been named nsP1a/4. The nonstructural N-terminal protein including nsP1a/1, nsP1a/2, and nsP1a/3 of the nsP1a polyprotein cannot interact with CD63, suggesting that nsP1a interacts with CD63 through its C-terminal region.

Tetraspanin proteins have been reported to be involved in the lifecycles of many viruses, such as human immunodeficiency virus (HIV)(37-39), hepatitis C virus (HCV) (40,41), human papillomavirus(HPV) (,42,43), human T cell lymphotrophic virus (HTLV) (44), porcine reproductive and respiratory syndrome virus (PPRSV) (45), and rotavirus(46). CD63 protein was confirmed to interact with the rotavirus VP6 protein and to be involved in the release of membrane vesicles by intestinal epithelial cells infected with rotavirus (47). CD63 also has been shown to play an essential role during HIV-1replication in macrophages. CD63 can interact with and accumulate in close proximity to the HIV gp41 protein during the cell-to-cell transfer of HIV (48).

We constructed CD63-knockdown and CD63-overexpressing cells lines to determine if CD63 expression affects HAstV-1 replication.CD63 overexpression increased the viral mRNA level in infected cells, whereas CD63 knockdown reduced the viral mRNA level in infected cells. A previous study suggested thatnsP1a/4 is involved inthe viral RNA replication process and mightbe necessary for efficient formation of the viral RNA-replication complex through a direct interaction with the viral RNA or with other proteins of the complex (14). Further research isneeded to investigate whether CD63 or nsP1a/4 is necessary forviral RNA or viral polymerase toform a functional replication complex and to identify which signal pathway isinvolved promoting virus replication.

We confirmed that CD63 is co-opted by HAstV nsP1a/4 during infection of susceptible host cells. The interaction between CD63 and nsP1a/4 appears to promote viral RNA replication. Our results confirmed that CD63 is involved in the formation of exosomes and is abundantly present in late endosomes and lysosomes. CD63 at the cell surface is endocytosed via a clathrin-dependent pathway. In late endosomes, CD63 is enriched on the intraluminal vesicles, which are secreted by specialized cells as exosomes via the fusion of endosomes with the plasma membrane. HAstV Yuc8 RNA replication occurs in association with host-cell membranes, and the entry of HAstV into host cells apparently depends on the maturation of endosomes (49).The significance of CD63 function in HAstV infection and whether CD63 is involved in HAstV cell binding, endocytosis, membrane vesicles, post-entry processing, or virus maturation still remains to be determined. We hypothesize that the CD63/nsP1a interaction plays a role in viral RNA replication, and further study is needed to verify that hypothesis.

## MATERIALS AND METHODS

### Cell culture and virus infection

HEK293T and Caco-2 cells were maintained in Dulbecco’s Modified Eagle’s Medium (DMEM; GIBCO, UK) containing 10% fetal calf serum (GIBCO, UK). A plasmid-based reverse genetics system for human astrovirus type 1 (HAstV-1) was provided by professor Akira Nakanishi, Department of Aging Intervention, National Center for Geriatrics and Gerontology Morioka, Obu,Aichi, Japan. An HAstV-1cDNA plasmid (pCAG-AVIC) that drives HAstV-1 cDNA expression from the CAG promoter was transfected into HEK293T cells. Cell-culture supernatants were collected 48 h after transfection and used as a source of recombinant viruses to infect Caco-2 cells (50). The virus was activated with 200 μg/mL porcine trypsin(Sigma) for 1h at 37°C before exposure to the cells. The virus titers in the Caco-2 cell cultures were determined by fluorescent-focus assays as described previously (50).

### Construction and expression of plasmids

HAstV-1 nsP1a protein (GenBank accession number: AGX15183.1) was amplified using primers listed in Table 1 and cloned into pGBKT7 vector (Clontech USA, Mountain View, CA) to generate BD-bait plasmid (pGBKT7-nsP1a) for the Y2H system. To generate N-terminally HA-tagged nsP1a andnsP1a/4 expression constructs, nsP1a or nsP1a/4 frames were amplified using the primers listed in Table1. We extracted the total RNA of HAstV type 1 (HAstV-1) from Caco-2 cells and amplified nsP1a by RT-PCR. The PCR products were cloned into PEF-HA PGK Hygrp vector to create PEF-HA-nsP1a and PEF-HA-nsP1a/4 plasmids via the in-fusion HD Cloning method (TaKaRa Bio,Dalian,China). The expression plasmids pGEX-6P-1-nsP1a andpGEX-6P-1-nsP1a/4 weregenerated viathe insertion of nsP1a or nsP1a/4 cDNA between the *Bam*H I and *Xh*o I sites of pGEX-6P-1. The expression plasmid pET-29a-CD63-LELwas generated via theinsertion ofCD63-LEL cDNAbetween the *Xh*o I and *Bam*H I sites of pET-29. To generate N-terminally Flag-tagged CD63(GenBank accession number: NM_001040034.1) or CD63-LEL expression constructs, CD63 or CD63-LEL PCR products were cloned into pcDNA3.1(+) vector to create pcDNA3.1-3flag-CD63 and pcDNA3.1-3flag-CD63-LEL via the in-fusion HD Cloning method.

**Table 1.**
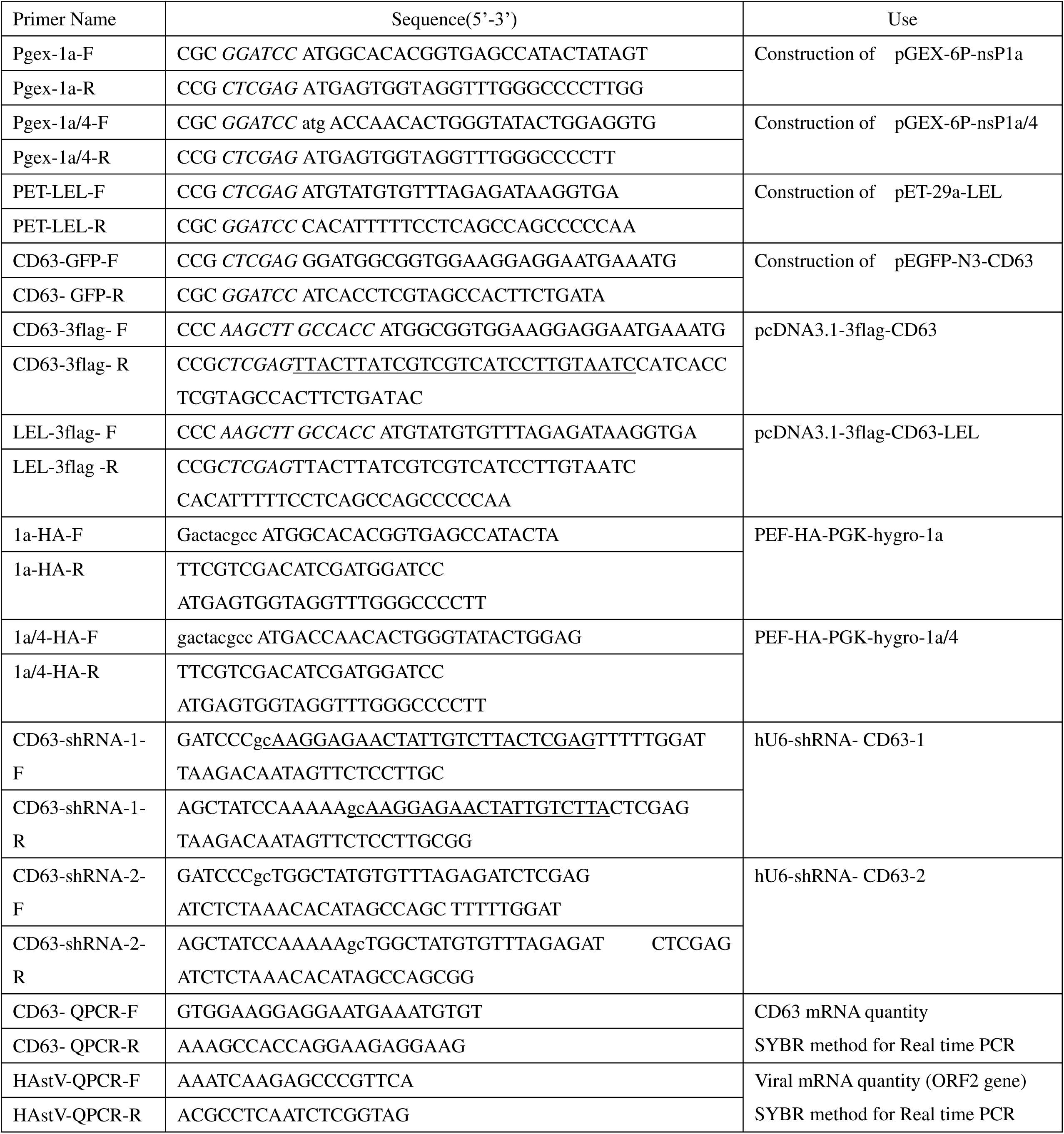
Oligonucleotides used in this study

### Expression and purification of recombinant nsP1a,nsP1a/4, and LEL proteins

The nsP1a and nsP1a/4 proteins were expressed and purified using thesame protocol was used for the CD63-LEL protein with some modifications. Briefly, the recombinant plasmid was transformed into competent *E. coli* BL21 (DE3) cells, which were then grown overnight in 5 mL Luria-Bertani broth. The incubation was continued until the optical density (OD600) reached 0.6 –0.8. The expression of the protein was induced with 0.1–1 mM IPTG at 25–37°C withshaking at 220 rpm. Expression of the protein was then analyzed by SDS-PAGE. Supernatant containing the fusion protein was purified using Glutahione Sepharose 4B (GE Healthcare Bio-Sciences, Little Chalfont, UK) according to the manufacturer’s instructions. The purification of recombinant nsP1a, nsP1a/4, and LEL proteins was further confirmed by western blot.

### Plasmid transfection

HEK293T cells were transfected with plasmids containing recombinant DNA using the Lipofectamine 2000 transfection reagent (Invitrogen) according to the manufacturer’s protocol. Briefly, cells were grown to approximately 90% confluence on six-well palates. The culture medium was replaced by serum-free medium containing the desired plasmid and Lipofectamine2000. The cells were incubated for 48 h at37°C in 5% CO_2_ and harvested for examination by western blot analysis to confirm the protein expression.

### Antibodies

Anti-Flag murine IgG1 monoclonal antibody, anti-HA murine IgG1 monoclonal antibody, rabbit antibody, anti-GST-Tag mouse polyclonal antibody, and anti-His-tag mouse polyclonal antibody were used (Sigma). Horseradish peroxidase-labeled goat anti-mouse IgG or goat anti-rabbit IgG (Santa Cruz Biotechnology Co.,Ltd, Shanghai, China) were used for western blotting. FITC-labeled goat anti-mouse IgG and Alexa Fluor^®^-labeled Goat Anti-rabbit IgG (Santa Cruz Biotechnology) were used as secondary antibodies for immunofluorescence and confocal microscopy. A polyclonal antibody to recombinant nsP1a or nsP1a/4 of HAstV was prepared in a previous study (51).

### Isolation of nsP1a-interacting cDNA clones using the Y2H technique

Afull-length expression cDNA library derived from the Caco-2 cell line was constructed for three ORFs. AY2H media set (Clontech,TaKaRa Biomedical Technology Co., Ltd.,Dalian, China) was used to identify nsP1a binding factors according to the manufacturer’s instructions. In brief, AH109 and Y187 cells were transformed with the bait (pGBKT7-nsP1a) and prey (Caco-2 cDNA library)constructs, respectively, according to the yeast transformation protocol (Clontech). A clone ofthe bait transformant was mated with a clone ofthe prey transformant and grown at 30°C overnight in 1 mL yeast extract peptone dextrose broth. The mated clones were selected on SD medium fortified with tryptophan and leucine to ensure successful mating. Finally, the interacting partners were screened on SD media lacking tryptophan (-Trp), leucine (-Leu), adenine (-Ade), and histidine (-His). Plasmids pGBKT7-53 and pGADT7-T, encoding the interacting protein pairP53 and Simian virus 40 (SV40) large T antigen, respectively, were used as positive controls. Plasmids pGBKT7-Lam and pGADT7-T, encoding the non-interacting protein pairlamin and SV40 large T antigen, respectively, served as negative controls.

### Western blot

Protein samples were separated by 12% SDS-PAGE and then transferred onto PVDF membranes (Millipore, Bedford, MA, USA). The membranes were incubated with anti-HA mAb(1:5000), anti-Flag mAb(1:10000), anti-CD63 mAb (1:2000), and anti-CD63 mAb (1:1000),respectively. The membranes were subsequently rinsed with PBST and treated with HRP-labeled goat anti-mouse IgG or goat anti-rabbit IgG as the secondary antibody (1:3000) (Santa Cruz Biotechnology). The proteins were visualized via scanning with ECL prime western blotting detection reagent (Beyotime Biotechnology,Beijing,China).

### GST pull-down assays

The recombinant pGEX-6P-1-nsP1a or pGEX-6P-1-nsP1a/4 plasmids were used to transform competent *E. coli* BL21 (DE3) cells. The cells were than grown overnight in5mL Luria-Bertani broth supplemented with ampicillin until the optical density (OD600) reached 0.6–0.8. Expression of pGEX-6P-nsP1a was induced with 0.1∼1mM IPTG at 25∼37°C with shaking at 220 rpm and then analyzed by SDS-PAGE. GST-nsP1a protein was purifiedusing a gravity-flow GST-Sefinose^™^Resin (Sangon Bitech, Shanghai, China) column according to the manufacturer’s instructions and detected by SDS-PAGE. His-tagged recombinant human CD63 protein was obtained from ThermoFisher Scientific Sino Biological (China).The expression and purification of the CD63 LEL domain were performed by the same method used for nsP1a and nsP1a/4. The Pierce™ GST Protein Interaction Pull-Down Kit (Thermo Fisher Scientific) was used to identify the interaction between nsP1a or nsP1a/4 and CD63 or CD63-LEL *in vitro*. In brief, prepared GST-nsP1a or GST-nsP1a/4 bait protein sample was bound to glutathione agarose. Then, the beads were washed four times with wash solution (25mMTris–HCl, 0.15MNaCl, pH 7.2). Pull-Down Lysis Buffer was incubated with recombinant His-tagged CD63 or His-CD63-LEL at 4°C for at least 1 h with gentle shaking. The eluted proteins were detected by SDS-PAGE followed by western blot analysis with anti-GST antibody and CD63 antibody.

### Co-immunoprecipitation of nsP1a/4 with CD63-LEL

HEK293T cells were transfected with PEF-HA-nsP1a/4 and pcDNA3.1-3flag-CD63-LEL expression plasmids. Following transfection for 48 h, the cells were lysed in NP-40 lysis buffer containing protease inhibitors. Following the manufacturer’s protocol, antibodies directed against Flag or HA were separately bound and cross-linked to Protein G Dynabeads (Novagen) using the reagents provided in the Pierce Crosslink Co-immunoprecipitation Kit (Pierce). Approximately 1–5 mg of untransfected or transfected cell lysates were then mixed with different sets of antibody-coupled beads and washed. The eluted proteins were analyzed by western blot using Flag or HA antibody.

### Immunofluorescence and confocal microscopy

HEK293T cells were co-transfected with PEF-HA-nsP1a/4 and pcDNA3.1-3flag-CD63-LEL plasmids. After 48 h, the cells were washed with 1 mL PBS and fixed with 4% paraformaldehyde for 10min at room temperature. After additional PBS washes, mouse anti-Flag and rabbit anti-HA was applied at a dilution of 1:1000.The cells were then incubated for 1h at room temperature, washed with PBS, and subsequently incubated with FITC-labeled goat anti-mouse IgG and Alexa Fluor^®^-labeled goat anti-rabbit IgG at a dilution of 1:500. The cells were then washed with PBS and observed under a confocal microscope.

### CD63 overexpression and knockdown

The primers used to construct pcDNA3.1-3flag-CD63 arelisted in Table 1. Two pairs of shRNAs targeting human CD63 and a negative control shRNA (Table 1) were cloned into hU6-MCS-CMV-GFP-SV40-Neomycin vector (Shanghai Genechem Co.,Ltd.) to generate CD63-shRNA-1, CD63-shRNA-2, and CD63-NC, respectively. HEK293T cells were transfected with the constructs along with pcDNA3.1-3flag-CD63 using Lipofectamine 2000 transfection reagent (Invitrogen, Carlsbad, CA) following the manufacturer’s protocol.

### Real-time RT-PCR

The target gene CD63 and HAstV RNA were quantified by real-time RT-PCR using the primers listed in Table 1. Total RNA was isolated from HEK293T cells using TRIzol (Invitrogen) according to the manufacturer’s instructions. cDNA was reverse transcribed from 1µg total RNA using PrimeScript Reverse Transcriptase (TaKaRa Bio,Dalian,China). Quantitative real-time RT-PCR was performed using theSYBR PrimeScript RT-kit(TaKaRa Bio) according to the manufacturer’s instructions.The relative transcript levels were analyzed using the ΔΔCt method. The results were analyzed using Bio-RAD IQ™5 optical system software.

### Statistical analysis

Data were presentedas the mean±SD. Differences betweengroups were examined using Student’s t test with P<0.05 as the threshold forstatistical significance.

## ACKNOWLEDGMENTS

This work was supported by grants from the National Nature Science Foundation of China (no. 81201285), the National Nature Science Foundation of Liaoning Province (no.20170540398), and the Biological Anthropology Innovation Team Project of JZMU (no.JYLJ201702)

